# Transmission and protection against re-infection in the ferret model with the SARS-CoV-2 USA-WA1/2020 reference isolate

**DOI:** 10.1101/2020.11.20.392381

**Authors:** Devanshi R. Patel, Cassandra J. Field, Kayla M. Septer, Derek G. Sim, Matthew J. Jones, Talia A. Heinly, Elizabeth A. McGraw, Troy C Sutton

## Abstract

SARS-CoV-2 has initiated a global pandemic and vaccines are being rapidly developed. Using the reference strain SARS-CoV-2 USA-WA1/2020, we evaluated modes of transmission and the ability of prior infection or vaccine-induced immunity to protect against infection in ferrets. Ferrets were semi-permissive to infection with the USA-WA1/2020 isolate. When transmission was assessed via the detection of vRNA at multiple timepoints, direct contact transmission was efficient to 3/3 and 3/4 contact animals in two respective studies, while respiratory transmission was poor to only 1/4 contact animals. To assess the durability of immunity, ferrets were re-challenged 28 or 56 days post-primary infection. Following viral challenge, no infectious virus was recovered in nasal wash samples. In addition, levels of vRNA in the nasal wash were several orders of magnitude lower than during primary infection, and vRNA was rapidly cleared. To determine if intramuscular vaccination protected ferrets against infection, ferrets were vaccinated using a prime-boost strategy with the S-protein receptor-binding domain formulated with an oil-in-water adjuvant. Upon viral challenge, none of the mock or vaccinated animals were protected against infection, and there were no significant differences in vRNA or infectious virus titers in the nasal wash. Combined these studies demonstrate that in ferrets direct contact is the predominant mode of transmission of the SARS-CoV-2 USA-WA1/2020 isolate and immunity to SARS-CoV-2 is maintained for at least 56 days. Our studies also indicate protection of the upper respiratory tract against SARS-CoV-2 will require vaccine strategies that mimic natural infection or induce site-specific immunity.

**Importance:** The SARS-CoV-2 USA-WA1/2020 strain is a CDC reference strain used by multiple research laboratories. Here, we show the predominant mode of transmission of this isolate in ferrets is by direct contact. We further demonstrate ferrets are protected against re-infection for at least 56 days even when levels of neutralizing antibodies are low or undetectable. Last, we show that when ferrets were vaccinated by the intramuscular route to induce antibodies against SARS-CoV-2, ferrets remain susceptible to infection of the upper respiratory tract. Collectively, these studies suggest protection of the upper respiratory tract will require vaccine approaches that mimic natural infection.

## Introduction

In December of 2019 a novel coronavirus, designated severe acute respiratory syndrome coronavirus 2 (SARS-CoV-2), crossed the species barrier and began to cause human infections in Wuhan, China (1). The virus was readily able to transmit from person-to-person and subsequently spread around the globe initiating a pandemic (2). SARS-CoV-2 infection of humans causes a range of clinical illness from asymptomatic infection to fever, shortness of breath, and cough, and the infection can progress to severe pneumonia with associated mortality (3, 4). The clinical manifestations of SARS-CoV-2 are referred to as coronavirus disease (COVID-19), and at the time of writing, SARS-CoV-2 has infected over 50 million people worldwide causing more than a 1.3 million deaths (5). Early during the pandemic, the mode of viral transmission was unclear; however, the growing consensus is large respiratory droplets are the predominant mode of transmission, with small droplets (sometimes referred to as “aerosol”) transmission also playing a role under certain environmental conditions (i.e., indoors with poor ventilation)(6).

To characterize SARS-CoV-2 and develop countermeasures, numerous virus isolates have been obtained from infected patients. One isolate, SARS-CoV-2 USA-WA1/2020, is widely used by the research community and was recovered from a patient that returned to Washington State, USA, from China in January 2020 (7). In parallel, several animal models of SARS-CoV-2 have been developed. These models include transgenic mice expressing the human ACE2 receptor (8, 9), hamsters (10, 11), cats (12–14), ferrets (14–16), and non-human primates (17–19). The ferret is a well-established model of influenza transmission and immunity, and influenza vaccine-efficacy in ferrets correlates with efficacy in humans (20–24). Importantly, several intramuscular administered SARS-CoV-2 vaccine candidates have been advanced to phase III clinical trials, and while these vaccines show significant promise, it remains unclear if these approaches will confer protection against infection (25–27).

In the ferret model of respiratory virus transmission, direct contact transmission is assessed by housing an infected donor animal with a naïve cage-mate, while respiratory transmission, involves housing an infected donor adjacent to a naïve animal (designated respiratory contact) in cages that permit airflow between the animals but prevent direct contact. Importantly, in this system, large (>5 um) and small droplet (<5 um) transmission cannot be distinguished, and the term respiratory transmission encompasses transmission by airborne particles of both sizes. Since the emergence of SARS-CoV-2, several research groups characterized transmission of different viral isolates in ferrets (14–16, 28, 29). These groups have reported transmission was efficient (i.e., to 100%) to direct contacts, with variable transmission efficiency to respiratory contacts (i.e., to 33-75% of contacts)(15, 16, 28). Therefore, as both contact and respiratory droplet transmission have not been evaluated for the reference strain USA-WA1/2020, we sought to separately evaluate these modes of transmission in ferrets. Moreover, given that ferrets are a highly suitable model of immune-mediated and vaccine-mediated protection against influenza, we evaluated if prior SARS-CoV-2 infection or intramuscular vaccination against the S-protein receptor-binding domain (RBD) would protect against infection. Collectively, we demonstrate that the USA-WA1/2020 isolate transmitted efficiently by direct contact in ferrets and that prior infection conferred protection against re-infection, while intramuscular vaccination did not protect against infection.

## Results

### Evaluation of contact and respiratory transmission of SARS-CoV-2

To determine the predominant mode of transmission of the SARS-CoV-2 USA-WA1/2020 isolate in ferrets, direct contact and respiratory droplet transmission were evaluated separately. Animals (n=8) were inoculated with 10^5^ TCID50 of SARS-CoV-2 in a 1 mL volume. This volume permits infection of both the upper and lower respiratory tract (30). Six hours after virus inoculation, half of the animals were co-housed with a naïve cage mate, while the remaining animals were introduced into respiratory transmission cages and paired with a naïve respiratory contact. The respiratory transmission cages are designed such that the ferrets are separated by two offset perforated barriers that are 2.5 cm apart. Air flows from the side of the infected animal towards the naïve contact. After evaluating direct contact and respiratory transmission, a second confirmatory direct contact transmission experiment (n=4 donors and n=4 direct contacts) was performed. For all experiments, nasal wash samples were collected every other day from both the infected donor and contact animals, and vRNA and infectious titers were determined by qRT-PCR and tissue culture infectious dose 50% (TCID50), respectively. All experimental animals were monitored daily for clinical signs, and body weight and temperatures were taken every other day.

For all experimental animals, no overt clinical signs, body weight loss, or elevated temperatures were observed (data not shown). Analyses of levels of vRNA and infectious virus in donor animals are shown in Figure 1. Figure 1 panel A-C and G-I display the initial direct contact and respiratory transmission experiment, respectively. Results for the confirmatory direct contact study are shown in Fig 1 panels D-F. In the initial direct contact and the respiratory transmission study, donor ferrets had high levels of vRNA in the nasal wash from days 1-7 post-infection with levels declining at later time points; however, recovery of infectious virus was variable. Infectious virus could not be recovered from all animals between days 1-7 post-infection, and viral titers were low, between 10^1^ to 10^3^ TCID50/mL at most time points.

**Figure 1.**
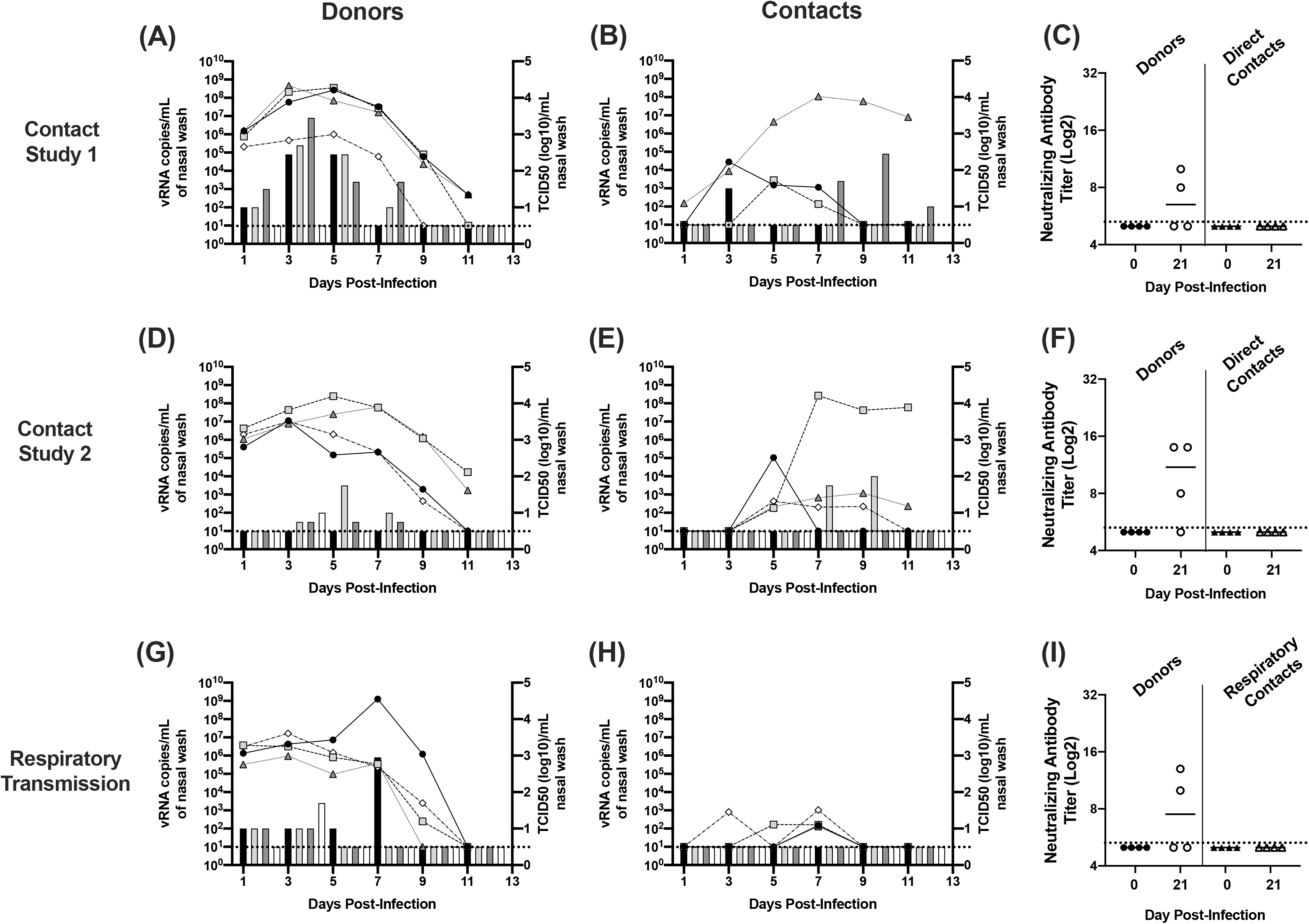
Direct contact and respiratory transmission of the SARS-CoV-2 USA-WA1/2020 isolate in ferrets. Panels A-C, D-F, and G-I display three separate transmission studies. Panels A-C, and D-F each represent a contact transmission study, while panels G-I display data from a respiratory transmission study. Panels A, D, G display nasal wash titers determined by qRT-PCR (left Y-axis) and TCID50 (right Y-axis) for the SARS-CoV-2 inoculated donor animals in each experiment. Line graphs indicate levels of vRNA and bar graphs indicated infectious titers. Panels B, E, and H similarly display nasal wash titers for contact animals. In a given panel, each shaded bar or symbol represents the same animal sampled over multiple time points. Paired donor and contact animals have the same shaded bar or symbol between panels. Panels C, F, and I show neutralizing antibody titers for each donor and contact animal. For all experiments, 4 pairs of ferrets (2 pairs of males and 2 pairs of females) were used, and nasal wash samples were collected every other day. Blood was collected on day 21 post-contact. In the first direct contact transmission study, panels A-C, one direct contact animal was removed due to fighting with its cage mate. Horizontal dashed line indicates limit of detection.

When viral shedding in direct contact ferrets was evaluated (Figure 1B), vRNA was detected in all of the contact ferrets at two or more time points. Infectious virus was recovered from one animal on day 3, and a second animal on days 7-11. One contact ferret was excluded from the study due to fighting with its cage mate. Thus, of the 3 direct contact ferrets, all 3 animals had vRNA in the nasal wash, and infectious virus was recovered from 2 of these animals. Analysis of the vRNA and infectious virus from the respiratory contact ferrets (Figure 1H) showed that vRNA could be detected at 1 or more timepoint in 3 of the 4 contacts; however, only one animal had vRNA present in the nasal wash at two consecutive time points. None of the animals displayed an increase in vRNA levels over time, and infectious virus could not be recovered from any of the respiratory contacts.

To verify our findings on contact transmission, we repeated the contact transmission experiment (Fig 1 D-F). Four donor animals were inoculated with the same dose of virus and each animal was housed with a naïve cage mate. As shown in Fig 1D, the donor animals had high levels of vRNA in their nasal wash, but again recovery of infectious virus was variable, and titers were low. When levels of vRNA were evaluated in the contact animals (Fig 1E), vRNA could be recovered from all 4 contacts at 1 or more time point, and three of the contacts had vRNA in the nasal wash at 3 or more time points. Infectious virus was only recovered from a single ferret on day 7 and 9 post-contact.

To determine if ferrets were seroconverting upon virus challenge and if any of the contact ferrets developed SARS-CoV-2 antibodies, microneutralization assays were performed on serum collected on day 21 post-contact. The levels of neutralizing antibodies are shown in Fig 1 panels C, F, and I. In all 3 transmission experiments, 2 or 3 of the 4 donor animals, and none of the direct contact or respiratory contact animals developed neutralizing antibodies. Combined these results indicate that ferrets are semi-permissive to infection with the SARS-CoV-2 USA-WA1/2020 isolate; however, assessment of transmission efficiency depends upon which assay is used to assess viral infection. If the presence of vRNA at any time point is used to assess infection, transmission is highly efficient to 7 of 7 direct contacts and 3 of 4 respiratory contacts. Alternatively, if the isolation of infectious virus or seroconversion are used to assess transmission, direct contact transmission is moderate or poor to 3 of 7 or 0 of 7 direct contacts, respectively. For respiratory transmission, assessment by detection of infectious virus or seroconversion is also poor to 0 of 4 respiratory contacts. Given neutralizing antibody titers were low or undetectable for donor animals and recovery of infectious virus was highly variable, neutralizing titers and recovery of infectious virus likely underestimate infection. Thus, to assess transmission efficiency, a suitable approach may be to use the detection of vRNA at two or more consecutive time points as this would indicate the persistence of vRNA in the nasal cavity for at least 3 days. Using these criteria, direct contact transmission is efficient to 6 of 7 contacts, and respiratory droplet transmission is inefficient to 1 of 4 animals.

### Protection against re-infection

To determine if prior infection was protective against re-infection, the donor ferrets from the initial respiratory and direct contact transmission studies were re-challenged at day 28 and 56 post-primary infection, respectively. After virus challenge, animals were monitored for clinical illness, and nasal wash samples were collected every other day for 9 days to assay for levels of vRNA and infectious virus.

Following secondary virus challenge, none of the animals displayed clinical signs, weight loss or elevated temperatures (data not shown). As shown in Fig 2C, 1 of 4 animals had neutralizing antibody titers at the time of challenge on day 28, and 2 of 4 animals had titers on day 56. The animals challenged on day 28 were the donors from the respiratory transmission study (Fig. 1G), and neutralizing titers declined to below the limit of detection in one animal between day 21 and 28. Upon virus challenge at day 28, all 4 animals had moderate levels of vRNA on day 1 and this was cleared by day 3 (Fig 2A). For animals challenged on day 56 (Fig 2B), all 4 animals similarly had vRNA in the nasal wash on day 1, and 2 animals had vRNA in the nasal wash on day 3. No vRNA was detected at time points later than day 3, and for both the day 28 and day 56 challenge, no infectious virus was recovered at any time points post-infection. Importantly, of the animals challenged on day 56, the two animals that had vRNA in the nasal wash on day 3 did not have neutralizing antibody titers at the time of challenge. The presence of vRNA at two consecutive time points indicates these animals may have been infected and maintained low levels of replicating virus for several days. However, comparison of the levels of vRNA during primary (Fig 1 A and G) and secondary challenge (Figure 2) shows that upon re-challenge vRNA levels were several logs lower and vRNA was rapidly cleared. Collectively, these results indicate ferrets are protected against re-challenge for at least 56 days post-primary infection.

**Figure 2.**
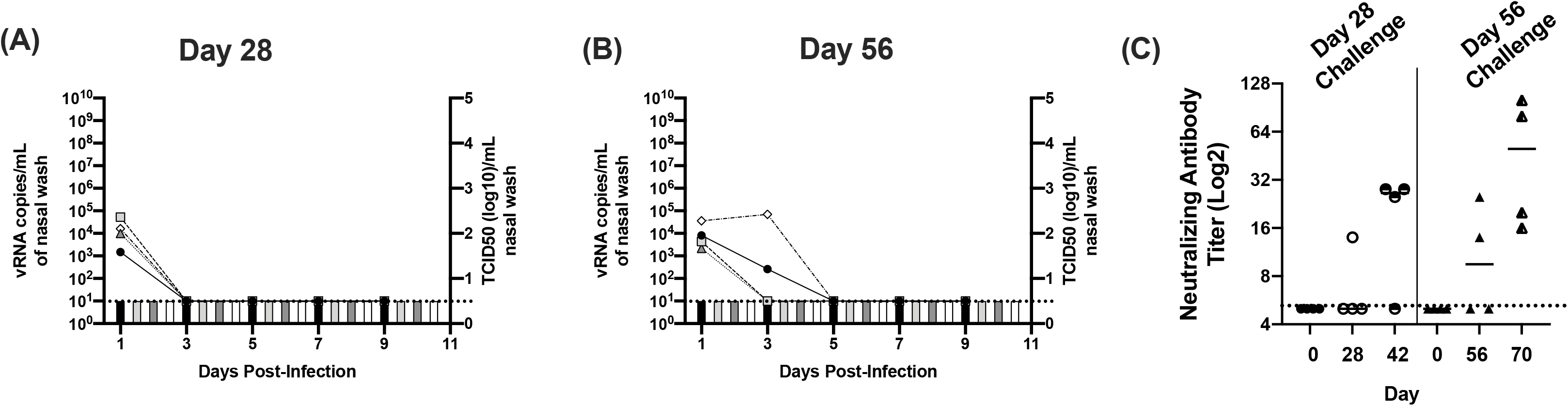
Viral and antibody titers in ferrets re-challenged with SARS-CoV-2 on day 28 and 56 post-primary infection. Panel A and B display nasal wash titers in ferrets re-challenged with SARS-CoV-2 on days 28 and 56 post-primary infection, respectively. Line graphs indicate levels of vRNA determined via N2 gene qRT-PCR (left Y-axis) and bar graphs indicated infectious titers (right Y-axis) determined via TCID50 on Vero cells. Panel C displays neutralizing antibody titers prior to primary infection (day 0), at the time of re-challenge (day 28 or 56) and 14 days post re-challenge (days 42 and 70). Horizontal dashed line indicates limit of detection.

### Intramuscular vaccination against the SARS-CoV-2 S-protein RBD does not protect against infection

Multiple intramuscular SARS-CoV-2 vaccine candidates are being advanced to clinical trials and these vaccines have been shown to induce neutralizing antibodies; however, it remains unclear if the induction of serum antibodies will reduce viral replication and/or prevent infection of the upper respiratory tract. Therefore, to determine if vaccine induced antibodies protect ferrets from SARS-CoV-2 infection, ferrets (n=4) received an intramuscular vaccine consisting of a potent oil-in-water adjuvant (Sigma Adjuvant System Adjuvant) alone (designated mock group) or the RBD of the SARS-CoV-2 S-protein mixed with adjuvant (designated RBD vaccine group) (31–33). Animals were given a primary vaccination and boosted on day 28. Every two weeks during the course of vaccination, blood samples were collected to assay for the presence of antibodies. On day 56 (i.e., 28 days post-boost vaccination) animals were challenged with SARS-CoV-2 and nasal wash samples were collected every other day for 13 days. As shown in Fig 3A and B, prior to virus challenge, none of the mock vaccinated animals developed binding or neutralizing antibodies against the RBD protein. In contrast, the vaccinated animals displayed increasing RBD binding antibodies over time with a boost in antibody titers following the secondary vaccination. At the time of virus challenge on day 56, the neutralizing activity of antibodies induced by vaccination was verified by microneutralization assay (Fig 3B) and titers ranged from 1:32 to 1:320.

**Figure 3.**
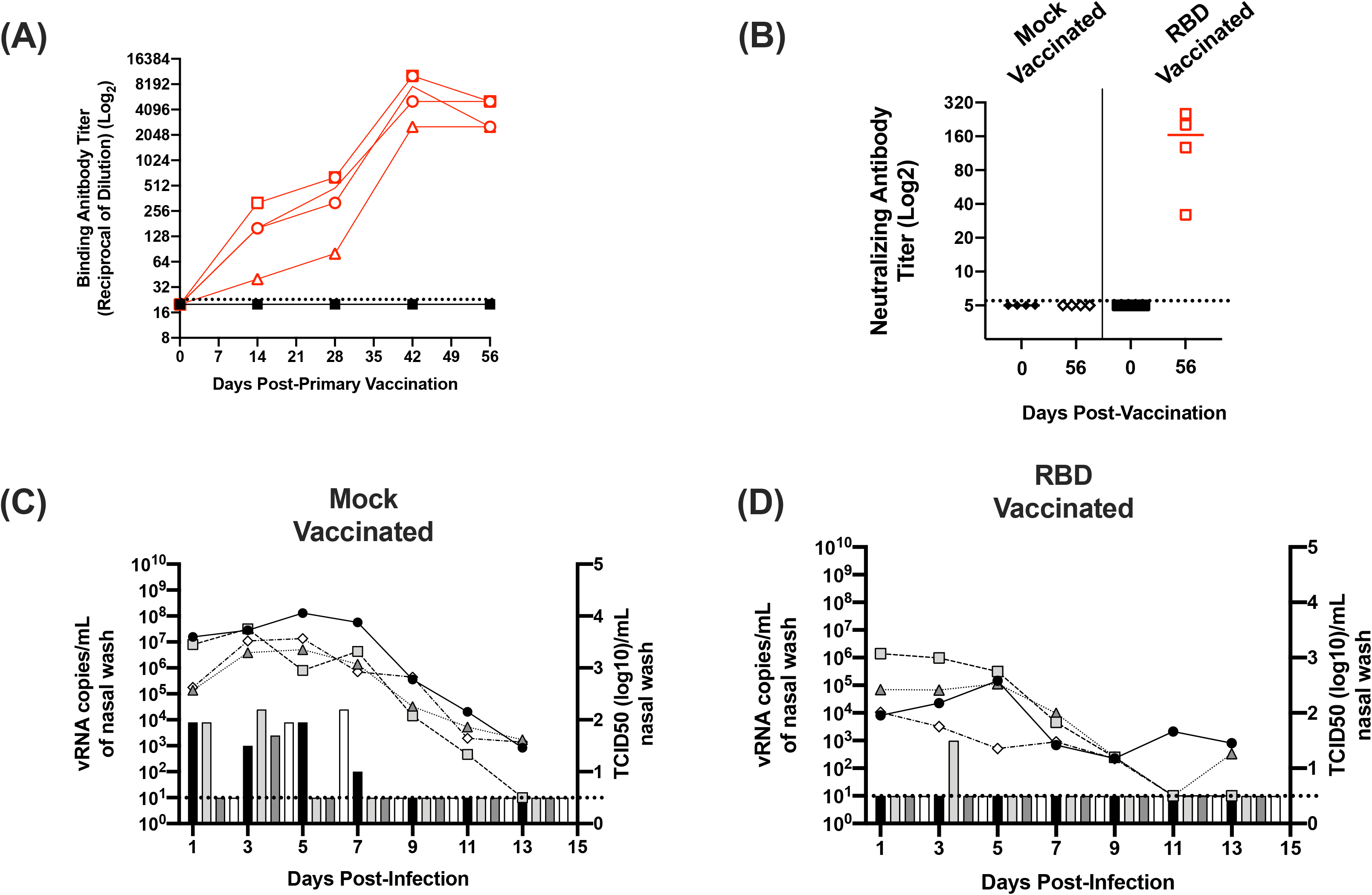
Antibody and viral titers in SARS-CoV-2 infected mock and RBD vaccinated ferrets. Panels A displays binding antibody titers against the S-protein RBD determined by ELISA on days 0, 14, 28, 42, and 56 post-primary vaccination. Red open symbols represent RBD vaccinated ferrets. Closed black symbols represent mock vaccinated animals. Animals were given a secondary vaccination on day 28. Panel B displays neutralizing antibody titers on day 56. Panel C and D display nasal wash titers in mock and RBD vaccinated animals challenged with SARS-CoV-2, respectively. Line graphs indicate levels of vRNA determined via N2 gene qRT-PCR (left Y-axis) and bar graphs indicated infectious titers (right Y-axis) determined via TCID50 on Vero cells. Horizontal dashed line indicates limit of detection.

On day 56, 28 days post-secondary vaccination, ferrets were challenged with 10^6^ TCID50 of SARS-CoV-2. For these studies a 10-fold higher challenge dose was utilized to enhance the recovery of infectious virus from the mock vaccinated animals. As shown in Fig 3C, following virus challenge, vRNA was detected in all of the mock vaccinated animals for at least 9 days. Levels of vRNA ranged from 10^3^ to 10^8^ copies per mL of nasal wash. As observed for the transmission studies, recovery of infectious virus was variable; however, all of the mock vaccinated animals shed infectious virus at one or more time point with titers between 10^2^ to 10^4.5^ TCID50/mL. In the RBD vaccinated animals, vRNA was similarly detected in the nasal wash samples for several days post-infection. Compared to the mock vaccinated animals, RBD vaccinated animals had ~10-100-fold lower titers of vRNA in the nasal wash between 10^2^ to 10^6^ copies per mL, and infectious virus was only recovered from a single animal on day 3. While both measures indicated a trend towards reduced viral load, there were no significant differences between the levels of vRNA or infectious virus between the mock and RBD vaccinated animals (*p*≥0.18). Combined these findings indicate vaccination against the RBD did not significantly reduce viral load in the nasal cavity or protect ferrets from infection with SARS-CoV-2.

## Discussion

Given the wide use of the SARS-CoV-2 USA-WA1/2020 strain in the research community, we evaluated transmission of this isolate in the ferret model and the role of pre-existing immunity induced via infection or vaccination on SARS-CoV-2 infection. When transmission by direct and respiratory contact were evaluated separately, the assessment of transmission efficiency depended on the stringency of the criteria used to evaluate infection in the contact ferrets. If we applied the human diagnostic standard of the presence of vRNA at any time point as evidence of infection, transmission was efficient to direct contact animals, with 3 of 3 and 4 of 4 contacts having vRNA in the nasal wash in separate studies, or a combined total of 7 of 7 contact animals being infected. If the recovery of infectious virus was used to assess transmission, transmission efficiency was reduced to 2 of 3 or 1 of 4 direct contacts in separate studies, or a total of 3 of 7 contacts. When seroconversion was evaluated, only a proportion of donor animals developed neutralizing antibodies and no contact animals seroconverted. In our respiratory transmission studies, the presence of vRNA was detected in 3 of 4 contacts, but infectious virus could not be recovered from any of the animals. When evaluated for seroconversion, 2 of 4 donor animals seroconverted, while none of the respiratory contacts developed SARS-CoV-2 antibodies.

Evaluation of the vRNA kinetics and recovery of infectious virus in the donor ferrets (Fig 1A, D, G) demonstrates, that despite high levels of vRNA, only low levels of infectious virus were recovered. Moreover, only a proportion of donor animals developed low titers of neutralizing antibodies, and none of the infected animals develop clinical signs or lost weight. Thus, the combination of poor virus recovery, limited seroconversion, and the absence of clinical signs indicate ferrets are semi-permissive to the SARS-CoV-2 USA-WA1/2020 isolate. Upon close evaluation of the vRNA levels in contact animals (Figure 1 B, E, H), we noted that some animals had vRNA present at a single time point while others had vRNA detected over multiple consecutive time points. Using the criteria of detection of vRNA at multiple consecutive time points as evidence of infection, 6 of 7 direct contacts and 1 of 4 respiratory contacts became infected.

Several other groups have reported on transmission of different SARS-CoV-2 isolates in ferrets (15, 16, 28, 34, 35). Our findings for the SARS-CoV-2 USA-WA1/2020 strain are most consistent with studies on the NMC-nCoV02 and Muc-IMB-1 strains (15, 16). When these isolates were evaluated for direct contact transmission in ferrets, the recovery of infectious virus was variable from both donors and contact animals; however, when the detection of vRNA alone was used to assay for transmission to direct contacts, both viruses were detected in 100% of contact animals (15, 16). Similar to our findings, when the NMC-nCoV02 strain was evaluated for airborne transmission, only viral RNA was detected in contact animals, and transmission occurred to 2/6 animals (15). Evaluation of neutralizing antibody titers in donor and direct contact animals for the Muc-IMB-1 isolate indicated that all 3 donors and 1 of 3 infected contacts seroconverted, while for the NMC-nCoV02 isolate all donors and direct contacts seroconverted, but only 1 of 2 positive respiratory contacts developed antibodies (16).

Both direct contact and respiratory transmission have also been evaluated for the BetaCov/Munich/BavPat1/2020 isolate (28, 35). For this virus, transmission was observed to 100% of direct contact animals determined by the presence of vRNA, recovery of infectious virus, and seroconversion. When respiratory transmission was evaluated, depending on the study design, transmission occurred to 2 of 4 or 3 of 4 contacts, and this was confirmed by all 3 measures of transmission: presence of vRNA, recovery of infectious virus, and seroconversion. Importantly, for all infected donor animals, high levels of vRNA combined with low infectious titers were reported, while infected direct or respiratory contacts had further reduced levels of vRNA and infectious titers (28, 35). A recent report has also evaluated direct contact transmission of the SARS-CoV-2 USA-WA1/2020 isolate (34). This study evaluated the efficacy of a nucleoside analog inhibitor (MK-4482) to inhibit SARS-CoV-2 transmission. In untreated animals, the SARS-CoV-2 USA-WA1/2020 isolate transmitted to all co-housed direct contacts, and both vRNA and infectious virus were recovered from the direct contacts. When donor ferrets were treated with the MK-4482, vRNA but no infectious virus was detected in the nasal wash, and none of the direct contact animals became infected (34). In these studies, all animals were euthanized 72 hours post-contact, therefore seroconversion could not be evaluated. In contrast to our findings, it is unclear why infectious virus could be consistently recovered in this study, while recovery of infectious virus was variable in both our inoculated donors and direct contacts. Interestingly, limited recovery of infectious virus does not appear to be a feature of only our studies. Several other groups have reported recovery of low and variable virus titers from ferrets (14–16). One possible explanation for these differences is passage history of the viruses used in multiple studies. Our SARS-CoV-2 USA-WA1/2020 isolate was obtained at passage 5 and passaged once in Vero E6 cells. As repetitive passaging of SARS-CoV-2 has been shown to alter the quasi species (36, 37), it is possible our passage 5 virus had limited quasi species diversity, while lower passage isolates used by other groups may have maintained more diversity, thus facilitating infection and/or recovery from ferrets.

After completing the transmission studies, we re-challenged subsets of the donor animals on day 28 and 56 and demonstrated protection against re-infection. Following virus challenge on days 28 and 56, infectious virus was not recovered from any animals. vRNA was present in the nasal wash of all animals on day 1 in both groups. In the animals challenged on day 28, vRNA was cleared by day 3, while in two of four animals challenged on day 56, vRNA was also present on day 3, and this was cleared by day 5. While vRNA was detected at two time points in the animals challenged on day 56, the levels of vRNA in the nasal washes were several orders of magnitude lower than during the primary infection indicating immune-mediated control of the virus. Our findings are consistent with other groups that show protection against re-infection 28 days post-infection in ferrets, hamsters, and non-human primates (11, 17, 29); however, we extend the window of protection to at least 56 days. Moreover, the observation that most animals had low or undetectable neutralizing antibodies titers prior to challenge indicates cellular immunity plays a significant role in protection against re-infection.

Last, we evaluated if the induction of high levels of serum antibodies via intramuscular vaccination would protect ferrets from infection. Ferrets were given a prime and boost vaccination 28 days apart consisting of the RBD protein formulated with an oil-in-water adjuvant. We specifically chose to vaccinate against the RBD because some vaccine approaches using the full-length SARS-CoV-1 S-protein potentiated vaccine-enhanced disease (38–41), while this did not occur when the RBD alone was used as a vaccine (42–47). Using this vaccine strategy, we induced high levels of antibodies (Fig 3C) measured by ELISA and confirmed by neutralization assay. Upon viral challenge, vRNA was detected in both the mock and RBD vaccinated animals for at least 9 days post-infection. Infectious virus was recovered from all of the mock vaccinated animals at one or more timepoints, while only a single RBD vaccinated animal shed virus; however, these differences were not significant, indicating that high levels of serum antibodies did not protect ferrets against infection. Given influenza vaccine efficacy in ferrets correlates with efficacy in humans, these findings indicate that an intramuscular vaccine may not prevent SARS-CoV-2 infection in humans. Our findings are consistent with SARS-CoV-2 vaccine studies in non-human primates (48-50). In these animals, regardless of the vaccine formulation, the induction of high levels of neutralizing antibodies conferred protection of the lower respiratory tract but did not protect against infection of the upper airways. It is important to note that ferrets did not display clinical disease and replication in the lower respiratory tract is not a feature of the ferret model of SARS-CoV-2 (14, 15). Therefore, our results do not suggest that current vaccines would not be effective at reducing severe disease. Indeed, licensed influenza vaccines which induce neutralizing antibodies in the blood, often reduce disease severity, but do not necessarily block infection (51, 52).

Collectively, we demonstrate ferrets are semi-permissive to infection with the SARS-CoV-2 USA-WA1/2020 isolate and that transmission of this isolate is efficient via direct contact and poor by respiratory droplet. We further demonstrate ferrets are protected against re-infection for at least 56 days. Using a vaccination strategy to induce serum neutralizing antibodies, ferrets were not protected against SARS-CoV-2 challenge. Thus, while initial human vaccines will likely reduce disease burden, to completely prevent SARS-CoV-2 infections, vaccine strategies that induce immunity in the upper respiratory tract comparable to natural infection will need to be developed.

## Materials and Methods

### Viruses and cells

The CDC reference strain SARS-CoV-2 USA-WA1/2020 was used for all studies. The strain was deposited by the Centers for Disease Control and Prevention and obtained through BEI Resources, NIAID, NIH: SARS-Related Coronavirus 2, Isolate USA-WA1/2020, NR-52281. The virus was obtained from BEI at passage 5 and cultured once in Vero E6 cells (ATCC) to generate a virus stock. Vero E6 cells were cultured in 10% FBS-DMEM (Hyclone) supplemented with non-essential amino acids (Corning), 2 mM L-glutamine, 1 mM sodium pyruvate, 1500 mg/L sodium bicarbonate, and antibiotics and antimycotics (Life Technologies) at 37°C in 5% CO_2_. SARS-CoV-2 was cultured in the equivalent media with the serum component reduced to 2% FBS. The virus stock was aliquoted and stored at −80°C until use. The titer of propagated virus stock was determined on Vero E6 cells in 24-well plates. Nasal wash samples were titrated using both 24 and 96 well plates. To enhance the detection of infectious virus, 100 uL of nasal wash sample was incubated on 2 wells of a 24-well plate for one hour. The nasal wash sample was removed and replaced with virus culture media. To assess peak titers in the nasal wash samples, 10-fold serial dilutions of virus were titrated in 96-well plates of Vero E6 cells. After inoculation onto Vero E6 cells, plates were incubated and scored 96 hours later for the presence of cytopathic effect (CPE). The tissue culture infectious dose 50% was calculated using the method of Reed and Muench (53).

To confirm the virus maintained the furin cleavage site and receptor-binding domain, viral RNA was extracted using a QIAmp vRNA mini kit (Qiagen), reverse transcribed using Superscript III Reverse Transcriptase with random hexamers (Life Technologies) and amplified using specific primers flanking each gene region with GoTaq DNA Polymerase (Promega). PCR products were run on an agarose gel, excised, and purified. Purified DNA was subject to Sanger sequencing. The resulting sequencing files were analyzed using Lasergene DNA star software and were confirmed to contain the furin cleavage site and receptor-binding domain sequence when compared to the published consensus sequence (GenBank: MT246667.1).

### Biocontainment and Animal Care and Use

All experiments with SARS-CoV-2 were conducted in the Eva Pell Laboratory for Advanced Biological Research which is the Pennsylvania State University’s biosafety level 3 enhanced laboratory. This facility is approved and inspected by the US Department of Agriculture (USDA) and Centers for Disease Control (CDC). All animal studies and procedures were conducted in compliance with relevant regulations and guidelines. All animal studies were approved by the PSU Animal Care and Use Committee under protocol no. 202001440. Equal number of male and female ferrets, 24-30 weeks old (Triple F Farms, Sayre, PA) were used for all studies. All animals were screened by hemagglutination inhibition assay and determined to be seronegative for currently circulating human influenza A viruses.

### Transmission Experiments

Ferret transmission experiments were performed using large stainless steel ventilated ferret cages (Allentown, NJ) modified to permit two ferrets to be separated by an offset perforated divider as described previously (54). For direct contact transmission experiments, ferrets were housed in the same caging system except the perforated divider was removed to allow the animals to share the same cage. To evaluate direct contact or respiratory transmission, donor ferrets (n=4) were sedated by intramuscular injection with a mixture of ketamine (30 mg/kg), xylazine (2 mg/kg), and atropine (0.05 mg/kg). Animals were then intranasally inoculated with 1 mL of virus culture media containing 10^5^ TCID50 of SARS-CoV-2. After virus inoculation animals were given the reversal agent atipamezole (0.5 mg/kg) via intramuscular injection and were housed separately in individual biocontainment ferret cages (Allentown, NJ). Six hours after virus inoculation, depending on the mode of transmission each ferret was paired with either a naïve direct contact or respiratory contact. The direct contact experiment was repeated for a total of 8 direct contact transmission pairs, while respiratory transmission was evaluated in 4 pairs of animals.

Starting 1-day post-viral inoculation and every other day for 11-13 days, animals were sedated by i.m. injection with a mixture of ketamine (20 mg/kg), xylazine (2 mg/kg), and atropine (0.05 mg/kg), and nasal wash samples were collected by instilling a 1 mL volume of PBS into the nostrils and inducing sneezing on a petri dish. An additional 1 mL volume of PBS was used to rinse the dish and the nasal washes samples were aliquoted and frozen at −80°C. All animals were monitored daily for clinical signs, and body weight and temperatures were collected at the time of nasal wash sample collection.

On day 21 post-viral inoculation and pair housing, the animals were deeply sedated, and blood was collected via cardiac puncture prior to euthanasia with an overdose of sodium pentobarbital. Eight of the donor animals were maintained for re-challenge studies. Blood was collected via the anterior vena cava from these animals, and the animals were given atipamezole and monitored until recovery. Blood samples were processed to recover serum and serum was stored at −80°C for evaluation of antibody titers.

### Re-challenge Experiments

To evaluate protection against re-infection, 8 donor ferrets from the transmission studies were maintained and blood samples were collected bi-weekly. Blood samples were collected from the anterior vena cava under sedation with ketamine, xylazine and atropine. On day 28 and day 56, four ferrets were re-challenged with 10^5^ TCID50 as described above. Following viral challenge, nasal wash samples were collected every other day for 9 days. On day 14 post-challenge, animals were sedated, blood was collected, and the animals were humanely euthanized.

### Vaccination and Challenge Studies

Ferrets were mock vaccinated (n=4) with Sigma Adjuvant System Adjuvant (SAS) alone or receptor-binding domain vaccinated (n=4) with 50 ug of recombinant SARS-CoV-2 S-protein RBD (SinoBiological) mixed with SAS. RBD was purchased as a lyophilized powder and resuspending in sterile PBS. Adjuvant was mixed 1:1 with the PBS for mock vaccinated animals or RBD protein solution, and at the time of vaccination, each animal received 500 uL of vaccine given as two 250 uL injections, one per hind leg. Animals were given a primary and secondary vaccination 28 days apart. On day 56 (i.e., 28 days post-boost), animals were sedated and intranasally inoculated with 10^6^ TCID50 of SARS-CoV-2 in a 1 mL volume. After viral challenge, nasal wash samples were collected every other day for 9 days, and animals were monitored for clinical signs and weight loss for 14 days.

### Neutralization Assay

The titers of neutralizing antibodies in ferret sera were evaluated by a microneutralization (MN) assay as previously described (55). Briefly, serial two-fold dilutions of heat-inactivated sera were prepared and mixed with 100 TCID50 of SARS-CoV-2 in an equal volume. The virus-serum mixture was incubated for one hour at room temperature and then overlaid in quadruplicate on Vero E6 cell monolayers in 96-well plates. Plates were incubated at 37°C for 4 days and scored for cytopathic effect. The neutralization titer was determined as the reciprocal of the serum dilution that neutralized virus as evidenced by the absence of CPE.

### S-protein RBD ELISA

To quantify anti-S protein RBD antibodies from ferret serum, 96-well ELISA plates (Nunc) were coated overnight with 2.0 ug/mL (100ng/well) of SARS-CoV-2 RBD S Protein (Sino Biological) in sodium bicarbonate buffer. Subsequently, plates were blocked for 2 hours at room temperature with PBST (0.05% Tween) with 3% goat serum and 5% skim milk powder. Plates were washed with wash buffer (i.e., PBST) and incubated for 2 hours at room temperature with serial 2-fold dilutions of heat-inactivated serum diluted in PBST with 1% milk powder. Plates were washed again and incubated for 2 hours at room temperature with HRP conjugated anti-ferret goat antibody (Rockland) diluted 1:10,000 in PBST and 1% skim milk powder. Following washes, OPD substrate solution (Sigma) was added to all wells and 3M HCl was added after 10 minutes to stop the reaction. The optical density was measured at 490 nm on SpectraMax iD3 plate reader (Molecular Devices) and an absorbance reading greater than three standard deviations above the mean day 0 value was considered positive.

### SARS-CoV-2 Quantitative Reverse-Transcription Polymerase Chain Reaction (qRT-PCR)

To assess levels of vRNA in nasal wash samples, RNA was extracted from 140uL of nasal wash using the QIAmp Viral RNA mini kit (Qiagen) according to the manufacture’s recommendations. RNA was eluted in 40 uL of buffer AVE. Quantification of levels of vRNA was performed according to the CDC protocol approved for Emergency Use Authorization (56). Purified RNA (5uL) was analyzed with the 2019-nCoV CDC qPCR N2 assay (IDT) using ‘qScript One-Step qRT-PCR, Low ROX (QuantaBio)’ master mix on an ABI 7500 Fast Real-Time PCR System (ABI).

### Statistical Analyses

For statistical analysis, nasal wash titers and levels of vRNA were compared between the mock and RBD vaccinated animals using a Mann-Whitney U test. Statistical analysis was performed with Prism GraphPad (v8.4.3) and *p<0.05* considered statistically significant.

## Acknowledgements

DRP, CJF, KMS, and TCS planned and performed all experiments with assistance from TAH and DGS. MJ performed the qRT-PCR analysis. DRP, CJF, KMS, and TCS drafted the manuscript. TCS and EAM revised and edited the manuscript. All authors have no competing interests. We would like to acknowledge the Animal Resource Program and the Pell Laboratory Facility Managers for assistance and support with animal studies. This research was supported by a seed grant from The Huck Institutes of Life Sciences at Pennsylvania State University, by NIAID Contract HHSN272201400004C (NIAID Centers of Excellence for Influenza Research and Surveillance, CEIRS) and USDA National Institute of Food and Agriculture, Hatch project 4605.

